# Bariatric surgery normalizes circulating glucocorticoid levels and lowers glucocorticoid action tissue-selectively in mice

**DOI:** 10.1101/2020.07.23.218404

**Authors:** Elina Akalestou, Livia Lopez-Noriega, Ioannis Christakis, Alexander D. Miras, Isabelle Leclerc, Guy A. Rutter

**Affiliations:** Section of Cell Biology and Functional Genomics, Imperial College London, Hammersmith Hospital Campus, Du Cane Road, W12 0NN London, United Kingdom; Section of Investigative Medicine, Division of Diabetes, Endocrinology and Metabolism, Department of Metabolism, Digestion and Reproduction, Imperial College London, Hammersmith Hospital Campus, Du Cane Road, W12 0NN London, United Kingdom; Endocrine and General Surgery, Nottingham University Hospitals NHS Trust, City Hospital campus, Hucknall Road, Nottingham, NG5 1PB Nottinghamshire, United Kingdom

**Keywords:** bariatric surgery, diabetes, adrenal glands, glucocorticoids, 11β-HSD11

## Abstract

**Background:** Glucocorticoids produced by the adrenal cortex are essential for the maintenance of metabolic homeostasis. Glucocorticoid activation is catalyzed by 11β-hydroxysteroid dehydrogenase 1 (11β-HSD1) and signalling is achieved through the glucocorticoid receptor (GR), a ligand-dependent transcription factor. Excess glucocorticoids are associated with insulin resistance and hyperglycaemia. A small number of studies have investigated the effects of bariatric surgery, a gastrointestinal procedure known to improve insulin sensitivity, on glucocorticoid metabolism, but the hypothesised mechanism is assumed to be via weight loss.

**Aim:** To investigate the effect of bariatric surgery on glucocorticoid metabolism in lean and obese mice.

**Methods:** Lean mice and HFD mice underwent Vertical Sleeve Gastrectomy (VSG) or sham surgery. Glucose and insulin tolerance tests were performed at four and ten weeks post operatively and circulating corticosterone was measured. Liver and adipose tissues were harvested from fed mice and 11β-HSD1 and GR levels were measured by quantitative RT-PCR or Western (immuno-) blotting, respectively.

**Results:** VSG did not cause excess weight loss in lean mice when compared to sham operated mice. However, both lean and HFD VSG mice displayed significantly improved glucose clearance and insulin sensitivity. Remarkably, VSG restores physiological corticosterone production in HFD mice and reduces11β-HSD1 levels at four and ten weeks post-surgery. Additionally, lean mice displayed significantly lowered mRNA levels of 11β-HSD1 in subcutaneous adipose tissue and GR in liver.

**Conclusions:** Bariatric surgery improves insulin sensitivity and reduces glucocorticoid activation at tissular level, under physiological and pathophysiological (obesity) conditions, irrespective of weight loss. The reduction of glucocorticoid exposure may represent an additional contribution to the health benefits of bariatric surgery. These findings point towards a physiologically relevant gut-adrenal axis.

## Introduction

Type 2 Diabetes (T2D) is associated with impaired insulin sensitivity in peripheral tissues and pancreatic β-cell dysfunction, which can lead to reduced insulin secretion (1). Despite an abundance of pharmacological, nutritional, exercise and behavioural interventions, T2D focus remains on disease *management*, rather than *remission* (2-5). Bariatric surgery, originally conceived as weight loss-aiding gastrointestinal surgery (6, 7), has been shown to cause long-term remission of T2D in many patients (8-10). These effects go beyond reductions in adipose tissue mass and weight loss, and numerous studies (11-13) have attempted to determine the exact pathways involved in order to replicate the bariatric effect in a less invasive way. These changes include substantial improvements in insulin sensitivity and hepatic gluconeogenesis alongside improvements in liver, cardiovascular, pancreatic islet and kidney function (14-17), pointing to a potential trans-organ communication axis. Importantly, the potential mechanisms investigated include possible changes in the secretion or action of multiple hormones (18, 19). One possibility that remains largely unexplored, however, is a ‘gut-adrenal’ axis, and specifically an altered role for glucocorticoids in regulating metabolism after bariatric surgery.

Glucocorticoids are produced by the adrenal cortex primarily under control of the hypothalamic-pituitary-adrenal (HPA) axis, However, it has recently became apparent that the fine-tuning and regulation of the adrenal system is also controlled by adrenocorticotropin-releasing hormone (ACTH)-independent mechanisms (20) including changes in growth factors, neuropeptides, cytokines and adipokines (20, 21). These include glucose regulating hormones such as insulin (22), glucagon (23, 24) and glucagon-like peptide 1 (GLP-1) (25, 26). Glucocorticoids are activated at tissue level by 11β-hydroxysteroid dehydrogenase 1 (11β-HSD11), which is present in most tissues (with the exception of pancreatic beta cells where this gene is “disallowed”) (27) and acts predominantly as an NADPH-dependent reductase to regenerate the active glucocorticoid receptor (GR) ligand cortisol (or corticosterone, in rodents) from inactive cortisone (28).

Although glucocorticoids are important for the maintenance of lipid homeostasis, excess glucocorticoids can result in an increase in the circulating free fatty acids and lipid accumulation in skeletal muscle and liver, both of which are associated with insulin resistance (29-31). Glucocorticoid metabolism is dysregulated in human obesity, where unbalanced cortisol levels and 11β-HSD11 activity are observed (32-34). Both GR and 11β-HSD11 overexpression in the liver and the adipose tissue have been linked to insulin resistance, dyslipidaemia and hypertension in rodents (35). Cortisol activation relies heavily on 11β-HSD1 and it has been suggested that the inhibition of this enzyme may be a key target for T2D and obesity treatments, especially to improve insulin sensitivity (36, 37).

Several observations have linked bariatric surgery to glucocorticoid metabolism, particularly in the context of tissular regulation of cortisol in obesity and post-operative weight loss (38, 39). Nonetheless, one study (40) showed that patients with obesity at one year post bariatric surgery displayed a significant reduction in adipose tissue 11β-HSD1 activity, when compared to lean non-operated controls. This observation points to an effect that may be caused by weight loss-independent mechanisms and may include the gastrointestinal tract manipulation itself as part of a gut-adrenal axis. One potential mechanism could be the effect of the widely reported post-operatively increased GLP-1 on glucocorticoid regulation (41-43). In the present study we aim to investigate the effect of Vertical Sleeve Gastrectomy (VSG), a commonly performed bariatric surgery procedure (44), on adrenal function under normal and pathophysiological and disease conditions, by using lean and HFD mice, respectively.

## Results

### Vertical Sleeve Gastrectomy improves glucose tolerance and restores insulin sensitivity

Lean VSG-treated mice experienced no significant weight loss four weeks post-surgery, when compared to sham operated mice (26.9 ± 2 vs. 28 ± 0.9 g) (Fig. 1A), yet they displayed improved insulin sensitivity and glucose tolerance (Fig. 1B, C). Specifically, VSG significantly enhanced glucose clearance following an intraperitoneal glucose injection (3g/kg) (p<0.01 at 15, 30, 60 and 90 min.) with an observed peak at 15 min. and glucose levels dropping to almost baseline concentration within 60 min (Fig. 1C). In contrast, in sham-operated mice, glucose peaked at 30 min. and did not fully recover within the first 90min of measurement. Although all lean mice were metabolically healthy at baseline, VSG-treated mice showed enhanced insulin sensitivity, as measured by circulating glucose concentration in response to an intraperitoneal insulin injection (Fig. 1B). The effects of VSG on glucose and insulin tolerance in hyperglycaemic HFD mice are described in (16). In brief, by post-operative week ten, diet-induced hyperglycaemic VSG-treated mice exhibited mild, non-significant weight loss when compared to sham mice (39.5 ± 1.6 vs. 47.1 ± 5.3 g) yet their glucose tolerance curve followed the same pattern described in the lean mice. Of note, the insulin sensitivity curves were also highly similar between VSG-treated lean and HFD mice as they both presented a continuous drop of glucose concentration up until 60 min.

**Figure 1:**
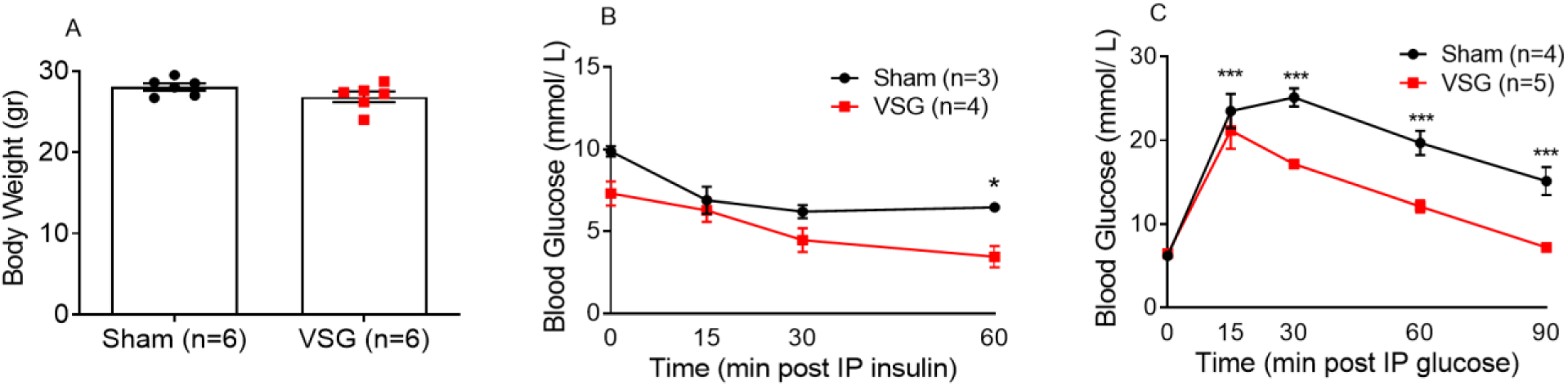
VSG improves glucose and insulin tolerance in lean and HFD mice. (A) Body weight at four weeks following VSG (n=6) or sham surgery (n=6) in lean animals. (B) Insulin tolerance test in lean mice. Human insulin (Actrapid, Novo Nordisk) was administered via intraperitoneal injection (0.8 IU/kg) after mice were fasted from 0900 to 1500 and blood glucose levels measured at 0, 15, 30 and 60 min. post injection, four weeks after surgery, n = 3-4 mice/group. (C) Glucose tolerance test in lean mice. Glucose was administered via intraperitoneal injection (3 g/kg) after mice were fasted overnight and blood glucose levels measured at 0, 15, 30, 60 and 90 min. post injection, four weeks after surgery, n = 3-5 mice/group. *P<0.05, ***P<0.001 VSG vs. Sham, following Student t-test or 2-way ANOVA. Data are expressed as means ± SEM.

### Vertical Sleeve Gastrectomy restores physiological corticosterone production in HFD mice

In order to assess the effect of VSG on glucocorticoid secretion, we measured corticosterone in blood collected from both lean and HFD mice that underwent VSG or sham surgery, at 0800 and 1900 within the same day. As expected in rodents, lean mice displayed low corticosterone levels during the morning (AM) and higher levels in the evening (PM), while no significant differences were observed between sham and VSG-treated mice. Interestingly, in HFD sham mice, corticosterone concentration remained at low levels both AM and PM, suggesting that the circadian rhythm of adrenal corticosterone is dysregulated by HFD. In contrast, in VSG-treated mice, PM corticosterone levels were restored to physiological levels observed in lean mice (Fig. 2A, B).

**Figure 2:**
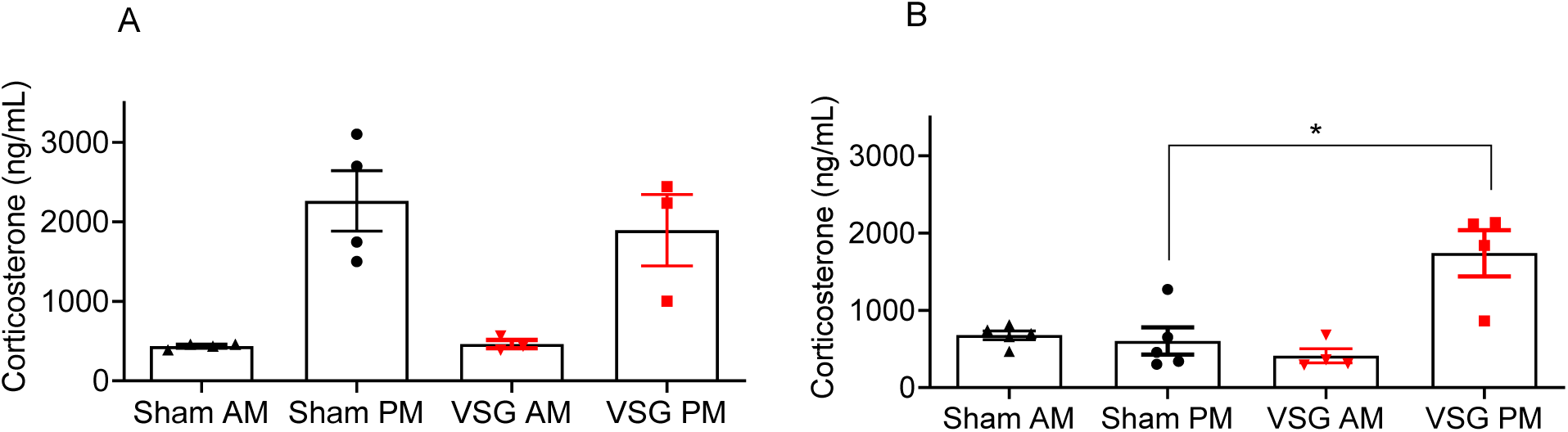
VSG restores physiological corticosterone production in HFD mice (A) Corticosterone measurement following blood collection at 0800 and 1900 within the same day in lean mice, four weeks post VSG or sham surgery (n=3-4). (B) Corticosterone measurement following blood collection at 0800 and 1900 within the same day in HFD mice, ten weeks post VSG or sham surgery (n=4). *P<0.05, VSG vs. Sham, following Student t-test. Data are expressed as means ± SEM.

### 11β-HSD11 is downregulated in liver and adipose tissue following Vertical Sleeve Gastrectomy

The metabolism of glucocorticoids on target tissues such as liver and adipose tissue are dependent on the enzyme 11β-HSD11. We therefore attempted to measure the expression of 11β-HSD11 in the liver and subcutaneous adipose tissue using biopsies from lean mice 4 weeks post-operatively, and HFD mice 10 weeks post-operatively. In both, lean and HFD mice, 11β-HSD11 mRNA levels were significantly reduced in the liver (Fig 3A, C). However, at the protein level, significant differences were only observed in lean mice (Fig 3B, G) and not HFD, although a tendency towards reduced protein levels could also be observed in HFD mice (Fig 3D, H).Moreover, 11β-HSD11 gene expression was significantly reduced in the adipose tissue of lean, but not in HFD mice (Fig 3E, F).

**Figure 3:**
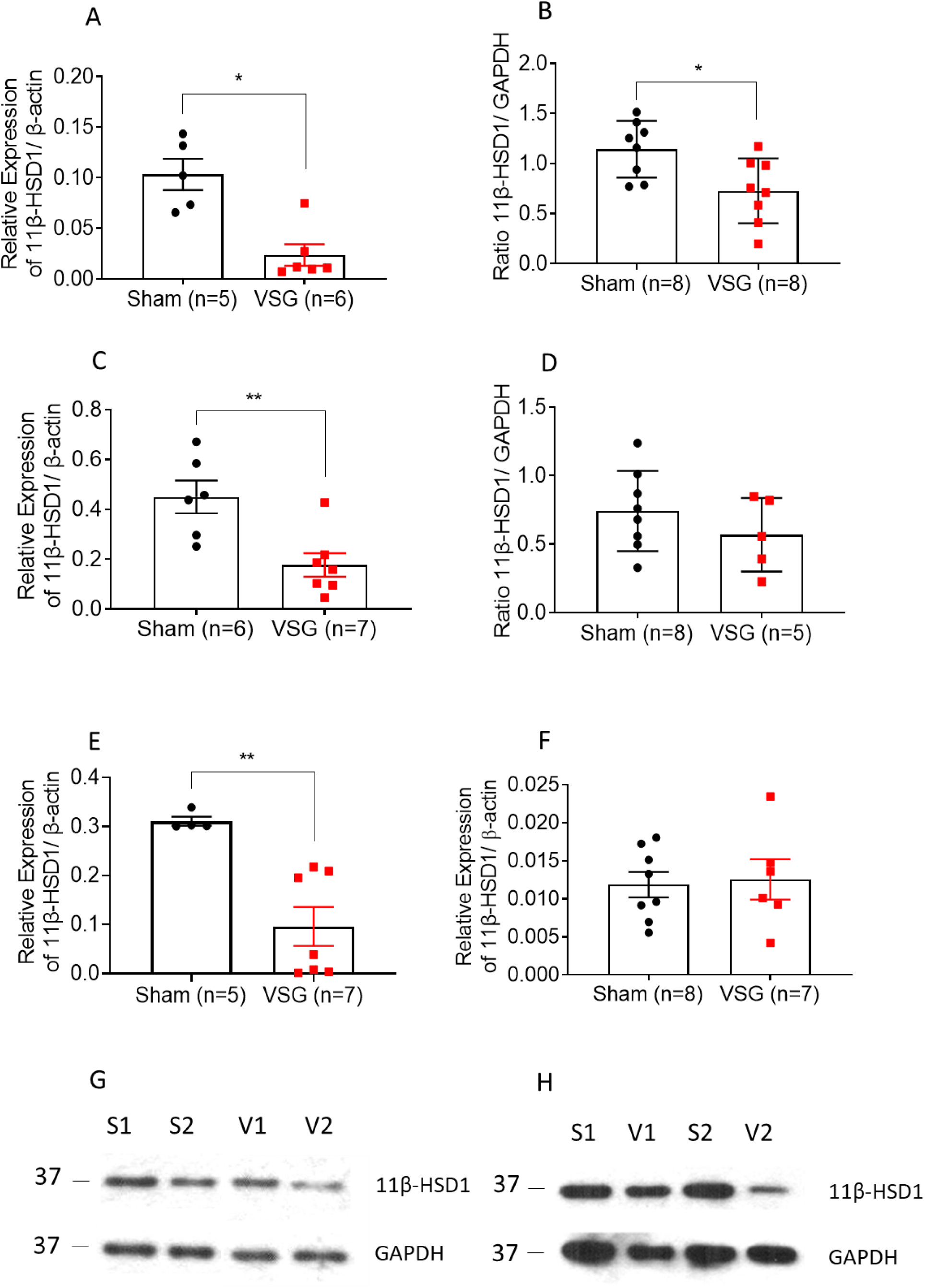
11β-HSD11 levels are decreased in the liver following VSG in mice (A) Quantitative PCR levels of 11β-HSD11 gene expression in liver of lean mice four weeks post VSG or sham surgery (B) Average intensity measurement from Western Blot immunoblotting quantification for 11β-HSD11 in lean mice four weeks post VSG or sham surgery (C) Quantitative PCR levels of 11β-HSD11 gene expression in liver of HFD mice ten weeks post VSG or sham surgery (D) Average intensity measurement from Western Blot immunoblotting quantification for 11β-HSD11 in HFD mice ten weeks post VSG or sham surgery. (E) Quantitative PCR levels of 11β-HSD1 gene expression in subcutaneous adipose tissue of lean mice four weeks post VSG or sham surgery (F) Quantitative PCR levels of 11β-HSD1 gene expression in subcutaneous adipose tissue of HFD mice ten weeks post VSG or sham surgery. (G) Example immunoblotting of lean sham and VSG mice liver using anti-rabbit 11β-HSD11 and anti-mouse GAPDH antibody (V1, 2 = VSG, S1, 2 = Sham) (H) Example immunoblotting of HFD sham and VSG HFD mice liver using anti-rabbit 11β-HSD11 and anti-mouse GAPDH antibody (V1, 2 = VSG, S1, 2 = Sham) *P<0.05, by Student’s t-test; **P<0.01 following Student t-test. Data are expressed as mean ± SEM.

### Lowered 11β-HSD11 expression reduction is not observed after incretin treatment

We next explored whether GLP-1 (the expression of which is reported to increase post bariatric surgery) (45) and/or lowered glycaemia, could play a role regulating the expression of 11β-HSD11. To this end, we injected the GLP-1 receptor agonist (GLP-1RA) Semaglutide subcutaneously at 5nmol/ kg in lean mice daily, for seven days. Although the body weight of the Semaglutide-treated mice did not change when compared to mice receiving saline (23.2 ± 1.7 vs. 23 ± 1.4 g) (Fig. 4A), fed glycemia was significantly lower on day seven (6.9 ± 0.5 vs. 12.5 ± 0.9 mmol/L) in the incretin vs saline-injected mice (Fig. 4B). However, the expression of 11β-HSD11 in liver and subcutaneous adipose tissue biopsies, obtained on the last day of treatment, showed no difference between Semaglutide and saline-treated groups, indicating that low glycemia or higher circulating GLP-1 does not replicate the effect observed following VSG (Fig. 4C, D).

**Figure 4:**
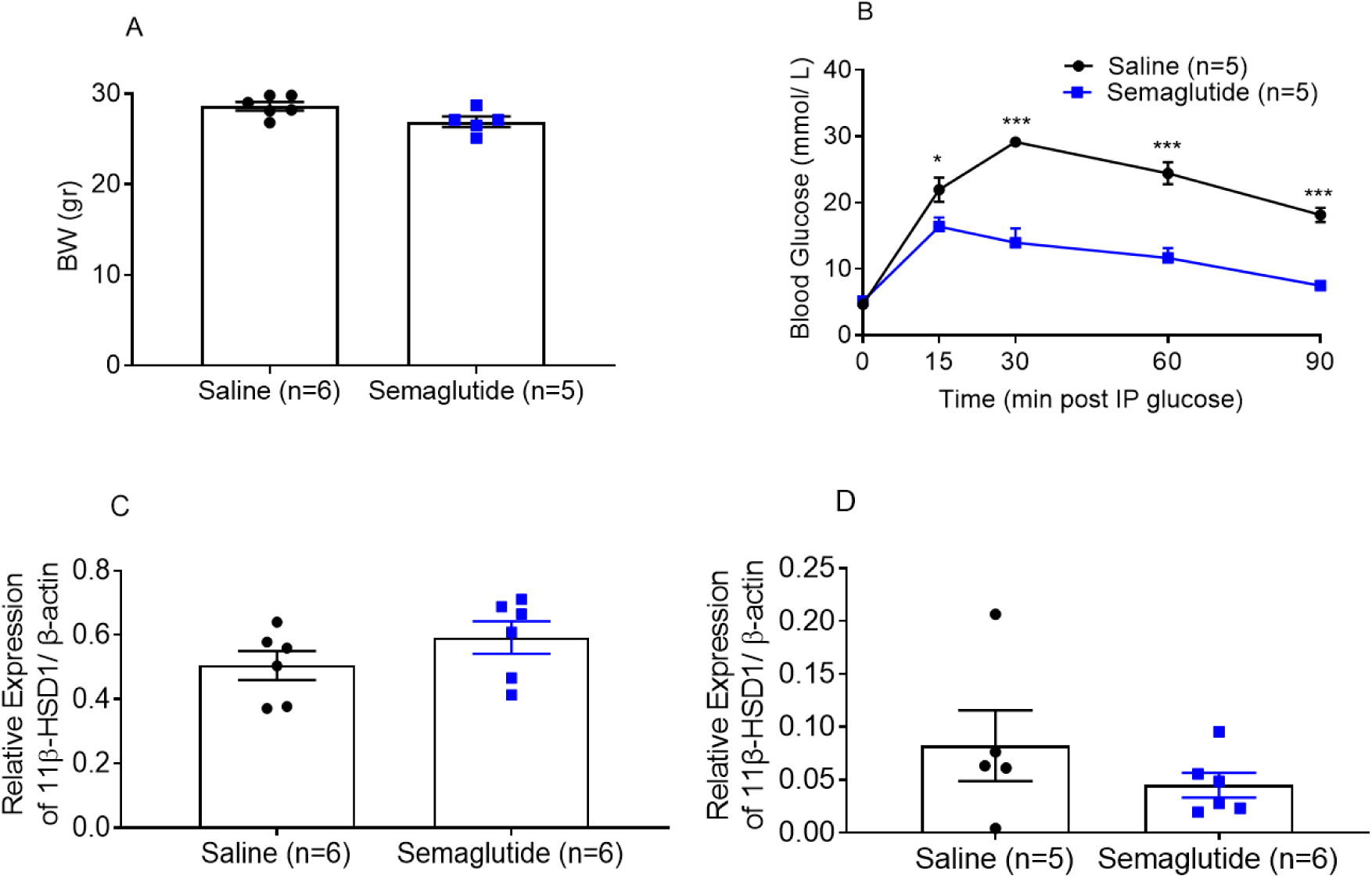
Semaglutide-induced glycaemia lowering does not decrease 11β-HSD11 levels significantly (A) Body weight measurement of lean mice that were treated with subcutaneous injection of 5nmol/kg semaglutide or saline for 7 days (B) Glucose tolerance test in lean mice following 7 days of semaglutide or saline injections. Glucose was administered via intraperitoneal injection (3 g/kg) after mice were fasted overnight and blood glucose levels measured at 0, 15, 30, 60 and 90 min. post injection (C) Quantitative PCR levels of 11β-HSD11 gene expression in the liver following 7 days of semaglutide or saline treated mice. (D) Quantitative PCR levels of 11β-HSD11 gene expression in the subcutaneous following 7 days of semaglutide or saline treated mice, n = 5-6 mice/group *P<0.05, ***<P0.001 by Student t-test. Data are expressed as means ± SEM.

### Glucocorticoid receptor is downregulated in the liver of lean mice following Vertical Sleeve Gastrectomy

Glucocorticoids act through the glucocorticoid receptor to regulate glucose metabolism in the liver, muscle, pancreas and adipose tissue, by controlling the expression of key enzymes. In light of our findings on 11β-HSD11 expression, we measured and compared GR gene expression in the liver of lean and HFD VSG-treated and Semaglutide-treated mice. GR receptor expression was significantly reduced in lean mice following VSG when compared to lean sham animals (Fig. 5A), although we did not observe any significant differences in the HFD (Fig. 5B). Importantly, lean mice treated with Semaglutide did not show any reduction in GR in the liver despite lower glycemia, corresponding to our previous findings on 11β-HSD11 expression (Fig. 3C).

**Figure 5:**
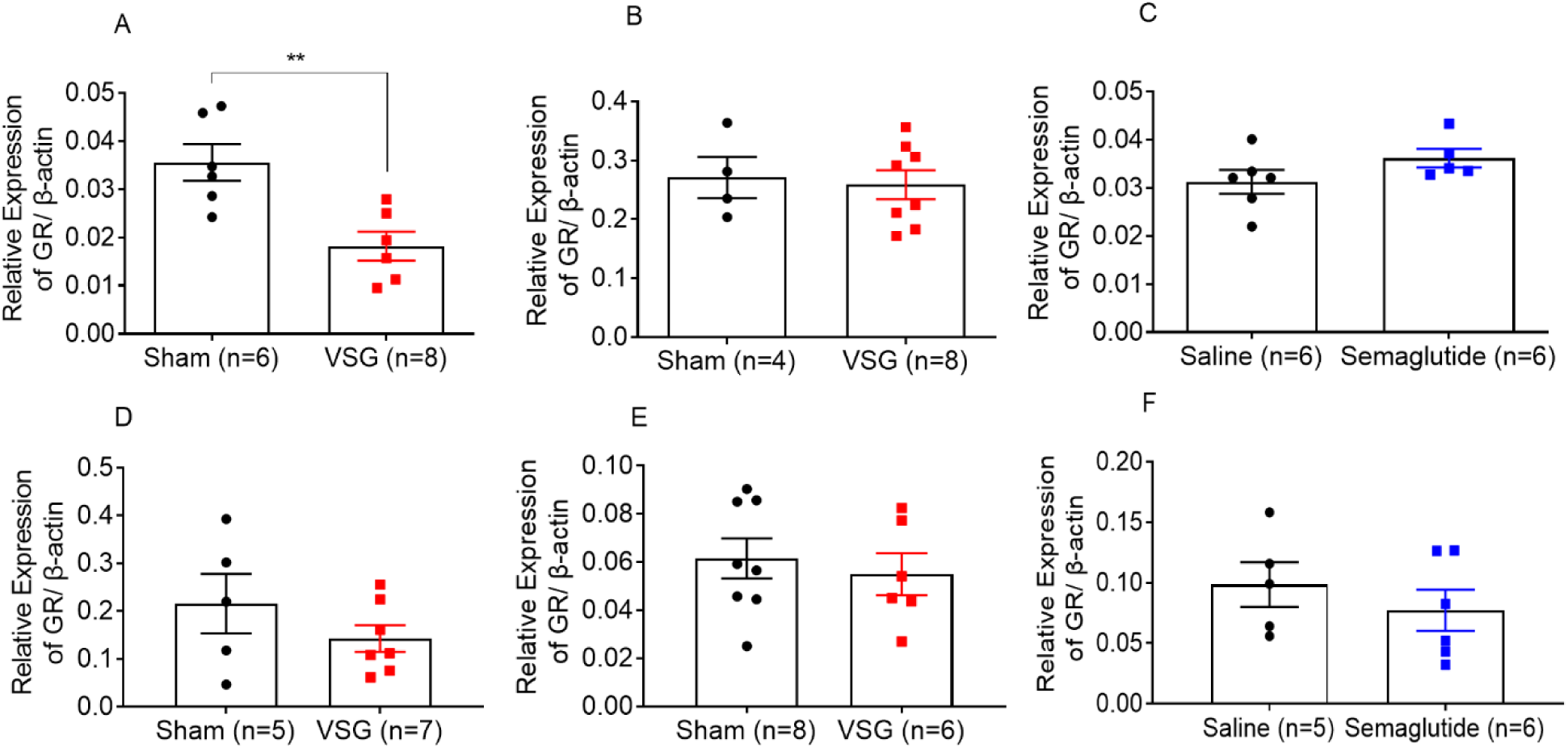
Glucocorticoid receptor levels in lean and HFD VSG or semaglutide-treated mice (A) Quantitative PCR levels of GR gene expression in liver of lean mice four weeks post VSG or sham surgery (B) Quantitative PCR levels of GR gene expression in liver of HFD mice ten weeks post VSG or sham surgery (C) Quantitative PCR levels of GR gene expression in liver of lean mice following 7 days of semaglutide (5nmol/kg) or saline (D) Quantitative PCR levels of GR gene expression in subcutaneous adipose tissue of lean mice four weeks post VSG or sham surgery (B) Quantitative PCR levels of GR gene expression in subcutaneous adipose tissue of HFD mice ten weeks post VSG or sham surgery (C) Quantitative PCR levels of GR gene expression in subcutaneous adipose tissue of lean mice following 7 days of semaglutide (5nmol/kg) or saline **P<0.001, by Student t-test. Data are expressed as means ± SEM.

## Discussion

In this study we explored, for the first time, the possibility of a gut-adrenal axis in physiology and pathophysiology. We utilised a gastrointestinal procedure that excludes most of the stomach and investigated the effects this would have on both circulating cortisol secretion and tissue-specific cortisol metabolism, while removing the confounding effects of weight loss. Furthermore, we attempted to identify the potential mediator of this axis by replicating the post-operative low glycemia pharmacologically by using Semaglutide.

The two most widely used types of bariatric surgery are currently VSG and Roux-n-Y-Gastric-Bypass (46). Despite the similar metabolic outcomes between the two procedures, VSG is reported to have lower mortality in mice (47, 48), but also allows for gradual weight regain, which is why it was chosen for this study (11, 49). Both lean and HFD-induced hyperglycemic models were deployed in order to compare glucocorticoid responses to bariatric surgery in models of health and disease. In the lean model, no weight loss was observed at four weeks post-operatively (Fig. 1A), yet glucose clearance increased significantly (Fig. 1C). In the HFD model, mild weight loss was observed at week ten post-operatively, yet glucose clearance rate increased significantly, as we have previously reported (16). These findings validate the metabolic phenotype of the VSG mouse model as reported in multiple studies (47, 50). Of note, insulin sensitivity curves between lean and HFD mice were almost identical (Fig. 1B) (16) demonstrating potentially similar mechanisms of improvement.

As an initial assessment of the glucocorticoid regulation following VSG, we measured circulating corticosterone in the early morning and late afternoon in all mice. This was done in four (lean mice) and ten (HFD mice) weeks after surgery to avoid measuring increased post-operative stress-related corticosterone levels. In healthy mice, physiological corticosterone levels are low in the morning and elevated in the evening (51), and both VSG and sham-treated lean mice demonstrated this pattern. Interestingly, HFD sham-treated mice showed no increase in corticosterone during the day, indicating impaired regulation of adrenal secretion. However, VSG appeared to restore normal corticosterone levels in HFD mice, matching the concentration found in lean mice, with a physiological increase in circulating glucocorticoid concentration apparent in the evening (Fig 2B). This is of importance as glucocorticoids, within the normal physiological range, are vital for metabolic, inflammatory and cardiovascular processes and an evening increase (equivalent of a morning increase in humans) could be associated with an overall improvement in lipid metabolism, immune response and vascular health (52). Moreover, the increase in corticosterone may contribute to improving functional enterocyte morphology and proliferation, and therefore gastrointestinal metabolic health in the long term, as cortisol is an important factor in the modulation of intestinal physiological functions. (53).

Despite the reported link between glucocorticoids and obesity (33), it is still unclear how circulating glucocorticoid levels are modulated following weight loss, and the cause of weight loss seems to be a confounder (54). Several studies have reported dysregulated cortisol levels in obesity, that remain unchanged, decreased, or increased following diet-induced weight loss, probably as a result of the stress associated with dieting, or the type of nutrients included in a diet (55-57). Several studies have attempted to study the regulation of cortisol levels following bariatric surgery in clinical settings (58-60). However, differences in the methodology and patient inclusion criteria (type of surgery, post-operative time, body mass index (BMI) or presence of T2D and/or eating disorders), have resulted in conflicting results. Our findings are supported by those of Valentine et al. who found a 54% rise in morning saliva cortisol levels at six and twelve months after VSG in women with obesity, but no differences in night-time samples, which corresponds with our findings (61).

Although the concentration of circulating glucocorticoids is important for their action, the effects of glucocorticoids on target tissues such as liver and adipose tissue are dependent on 11β-HSD1 activity (28). In the liver, glucocorticoids have been shown to stimulate gluconeogenesis by activating PEPCK and glucose-6-phosphatase (G6Pase) through the GR (62), while in the adipose tissue they stimulate lipolysis, resulting in the generation of glycerol to be utilized in gluconeogenesis and free fatty acids to be oxidised and used as energy source (63). Previous studies have shown that mice overexpressing 11β-HSD1 in adipose tissue develop visceral obesity, insulin resistance, dyslipidaemia, and hypertension (35), while liver-specific 11β-HSD1 overexpression results in insulin resistance and hypertension, but not obesity (64). In our study we found that 11β-HSD1 is significantly inhibited in the liver in both lean and HFD VSG-treated mice when compared to sham groups, pointing at reduced hepatic glucocorticoid activity (Fig 3. A, B, C, D) which can potentially be a contributing factor to the significant increase in insulin sensitivity observed on Fig 1B and E. Of note, 11β-HSD1 in the adipose tissue was only inhibited in the lean mice (Fig 3, E, F).

In an effort to elucidate the mechanism(s) that links VSG, a gastric procedure, to adrenal function, we explored the role of GLP-1, a post-prandially released hormone that enhances insulin secretion and lowers glucose concentration (65). This incretin hormone has also been shown to increase significantly following VSG, and to regulate glucocorticoid secretion (25, 26). We chronically injected a low dose of the GLP-1RA Semaglutide, to avoid significant body weight loss while achieving significantly lower glycemia, in lean mice as previously reported (17). Interestingly, 11β-HSD1 expression was not affected in either the liver or in the adipose tissue (Fig 4C, D) of Semaglutide-treated mice, suggesting that the observed effects are not driven by the reduction in blood glucose or a direct action of GLP-1.

Another key glucocorticoid function mediator is the GR, a nuclear receptor that, once bound to a glucocorticoid, translocates into the nucleus and regulates the expression of target genes (66). In the liver, GR has been shown to upregulate gluconeogenic enzymes (62) and its inactivation ameliorates hyperglycemia in diabetic mice (67). However, the GR role in the adipose tissue remains unclear as it has been found to play an important role in lipolysis and insulin resistance caused by exogenous glucocorticoids, but not insulin resistance caused by high fat feeding (68). Our study demonstrated that GR is downregulated in the liver in lean mice following VSG, but not following Semaglutide treatment or in HFD mice (Fig 5A, C). The GR expression in the adipose tissue was not altered in any of the mouse groups.

Although evidence suggest that bariatric surgery can affect the HPA axis (58, 69), there is more controversy around the direction of this regulation. The long-term effects of bariatric surgery on cortisol levels are also unclear. In clinical settings, hepatic 11β-HSD1 activity is primarily measured by calculating serum cortisol/cortisone ratio, but direct tissular expression requires biopsies which are usually not available for the liver. In patients with severe obesity following RYGB, 11β-HSD1 activity was increased in the liver but expression was decreased in the subcutaneous adipose tissue (38, 39, 70) highlighting that regulation of cortisol metabolism in obesity and after weight loss is highly tissue specific. In our study, we have demonstrated that, although VSG improves insulin sensitivity and inhibits 11β-HSD1 expression in the liver in both lean and HFD mice, the regulation of glucocorticoid metabolism in adipose tissue is different in both physiological and pathophysiological conditions.

### Limitations of the study

As with clinical studies, chronic cortisol measurements are required to further validate our findings. Moreover, studies on the exact transcription factors affected by the downregulation of 11β-HSD1 and GR are required to understand the pathway that links glucocorticoid metabolism to post-bariatric T2D remission. Similar to the Semaglutide study, more factors must be explored to pin a potential hormonal mediator between the gut and the adrenal secretion regulation. Finally, our results must extend to the clinical settings, especially in patients with cortisol oversecreting Cushing’s disease following bariatric surgery.

### Conclusions

We show here for the first time that VSG maintains normal corticosterone levels but inhibits tissular glucocorticoid metabolism in lean animals, by inhibiting 11β-HSD1 and GR, and that this effect is likely to be weight loss and glycaemia-independent. Moreover, we have demonstrated that corticosterone circulation is restored in obese VSG animals, while hepatic glucocorticoid metabolism is inhibited, even after weight regain. Insulin sensitivity and glucose clearance were enhanced in all groups. These observations demonstrate a potential new mechanism of T2DM remission after bariatric surgery.

## Materials and Methods

Animals -All animal procedures undertaken were approved by the British Home Office under the UK Animal (Scientific Procedures) Act 1986 (Project License PPL PA03F7F07 to I.L.) with approval from the local ethical committee (Animal Welfare and Ethics Review Board, AWERB), at the Central Biological Services (CBS) unit at the Hammersmith Campus of Imperial College London. Adult male C57BL/6J mice (Envigo, Huntingdon U.K.) were maintained under controlled temperature (21-23°C) and light (12:12 hr light-dark schedule, lights on at 0700). The animals were fed either PMI Nutrition International Certified Rodent Chow No. 5CR4 (Research Diet, New Brunswick, NJ) or 58 kcal% Fat and Sucrose diet (D12331, Research Diet, New Brunswick, NJ) for twelve weeks and ad libitum. Animals were exposed to liquid diet (20% dextrose) three days prior to surgery and remained on this diet for up to four days post operatively. Following this, mice were returned to either PMI or high fat/high sucrose diet. All mice were divided in two groups, VSG (n=7) and sham (n=6), and were euthanized, and tissues harvested, twelve weeks after surgery. Liver, adipose tissue and intestinal biopsies were removed from all mice at twelve weeks following sham or VSG surgery in the fed state, and were either snap frozen in −80°C, fixed in formalin, or both.

### Vertical Sleeve Gastrectomy

Anaesthesia was induced and maintained with isoflurane (1.5-2%). A laparotomy incision was made, and the stomach was isolated outside the abdominal cavity. A simple continuous pattern of suture extending through the gastric wall and along both gastric walls was placed to ensure the main blood vessels were contained. Approximately 60% of the stomach was removed, leaving a tubular remnant. The edges of the stomach were inverted and closed by placing two serosae only sutures, using Lembert pattern (71). The initial full thickness suture was subsequently removed. Sham surgeries were performed by isolating the stomach and performing a 1 mm gastrotomy on the gastric wall of the fundus. All animals received a five-day course of SC antibiotic injections (Ciprofloxacin 0.1mg/kg).

### Glucose Tolerance Tests

Mice were fasted overnight (total 16 h) and given free access to water. At 0900, glucose (3 g/kg body weight) was administered via intraperitoneal injection. Blood was sampled from the tail vein at 0, 5, 15, 30, 60 and 90 min. after glucose administration. Blood glucose was measured with an automatic glucometer (Accuchek; Roche, Burgess Hill, UK).

### Insulin Tolerance Tests

Mice were fasted for 8 h and given free access to water. At 1500, human insulin (Actrapid, Novo Nordisk) (0.8-1.5U/kg body weight) was administered via intraperitoneal injection. Blood was sampled from the tail vein at 0, 15, 30, 60 and 90 min after insulin administration. Blood glucose was measured with an automatic glucometer (Accuchek; Roche, Burgess Hill, UK).

### Plasma corticosterone measurement

To quantify circulating corticosterone levels, 50μl of blood was collected from the tail vein into heparin-coated tubes (Sarstedt, Beaumont Leys, UK) at 0800 and 1900. Plasma was separated by sedimentation at 10,000 ***g*** for 10 min. (4°C). Plasma corticosterone levels were measured in 10μl aliquots ELISA kits from Crystal Chem (USA).

### Western (immuno-) blotting

Liver and adipose tissue were lysed in ice-cold RIPA buffer containing a protease inhibitor mixture (Roche) and phosphatase inhibitors (Sigma-Aldrich). Lysates were denatured for 5 min at 95 °C in Laemmli buffer, resolved by 10% SDS-PAGE, and transferred to polyvinylidene difluoride membranes before immunoblotting. The following antibodies were used: anti-rabbit 11β-HSD1 (abcam, USA) (1:200) and anti-mouse GAPDH (Sigma-Aldrich) (1:1000). Intensities were quantified using ImageJ.

### RNA extraction, cDNA synthesis and Quantitative Polymerase Chain Reaction

Tissues were harvested and snap-frozen in liquid nitrogen. RNA was purified using PureLink RNA kit (Thermo Fisher Scientific, UK). The purified RNA was dissolved in RNase and DNase free distilled water (Thermo Fisher Scientific, UK) and was immediately stored at −80°C until further analysis. Complementary DNA was synthesized from total RNA with High-Capacity cDNA Reverse Transcription Kit (Thermo Fisher Scientific, UK), according to the protocol recommended by the manufacturer. Quantitative Reverse Transcription PCR (qRT-PCR) analysis was used to quantify the expression level of 11β-HSD11 and NR3C1 (GR) mRNA in kidney cortex, and adiponectin in SC adipose tissue. Primers, which crossed a splice junction, were designed using Primer Express (Invitrogen, UK; Table 1). The expression levels were measured by Q-PCR, using Fast SYBR Green Master Mix (Invitrogen) and a 7500 Fast Real-Time PCR System (Applied Biosystems, UK). Data from 11β-HSD11 and NR3C1 (GR) were normalized against β-actin levels. The analytical method used was 2(-Delta Ct).

**Table 1:**
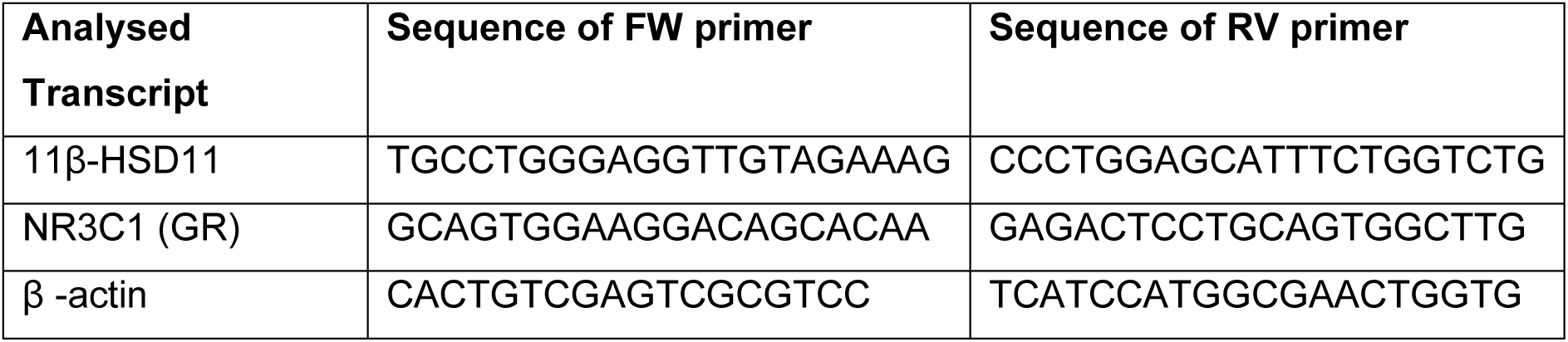
Primers Used for the Quantitative Detection of 11β-HSD11 and GR, normalised to β-actin.

### Statistical Analysis

Data were analysed using GraphPad PRISM 7.0 software. Significance was tested using unpaired non-parametric Student’s two-tailed t-tests with Bonferroni post-tests for multiple comparisons, or two-way ANOVA repeated measurements as indicated. P<0.05 was considered significant. Data are represented as the mean ± SEM.

## References

1. Kahn SE, Zraika S, Utzschneider KM, Hull RL. The beta cell lesion in type 2 diabetes: there has to be a primary functional abnormality. Diabetologia. 2009;52(6):1003–12.

2. Wycherley TP, Noakes M, Clifton PM, Cleanthous X, Keogh JB, Brinkworth GD. A high-protein diet with resistance exercise training improves weight loss and body composition in overweight and obese patients with type 2 diabetes. Diabetes Care. 2010;33(5):969–76.

3. Post RE, Mainous AG, 3rd, King DE, Simpson KN. Dietary fiber for the treatment of type 2 diabetes mellitus: a meta-analysis. J Am Board Fam Med. 2012;25(1):16–23.

4. Raz I, Hanefeld M, Xu L, Caria C, Williams-Herman D, Khatami H, et al. Efficacy and safety of the dipeptidyl peptidase-4 inhibitor sitagliptin as monotherapy in patients with type 2 diabetes mellitus. Diabetologia. 2006;49(11):2564–71.

5. Amori RE, Lau J, Pittas AG. Efficacy and safety of incretin therapy in type 2 diabetes: systematic review and meta-analysis. JAMA. 2007;298(2):194–206.

6. Sugerman HJ, Kellum JM, Engle KM, Wolfe L, Starkey JV, Birkenhauer R, et al. Gastric bypass for treating severe obesity. Am J Clin Nutr. 1992;55(2 Suppl):560S–6S.

7. Pories WJ, Swanson MS, MacDonald KG, Long SB, Morris PG, Brown BM, et al. Who would have thought it? An operation proves to be the most effective therapy for adult-onset diabetes mellitus. Ann Surg. 1995;222(3):339-50; discussion 50-2.

8. Mingrone G, Panunzi S, De Gaetano A, Guidone C, Iaconelli A, Nanni G, et al. Bariatric-metabolic surgery versus conventional medical treatment in obese patients with type 2 diabetes: 5 year follow-up of an open-label, single-centre, randomised controlled trial. Lancet. 2015;386(9997):964–73.

9. Courcoulas AP, Belle SH, Neiberg RH, Pierson SK, Eagleton JK, Kalarchian MA, et al. Three-Year Outcomes of Bariatric Surgery vs Lifestyle Intervention for Type 2 Diabetes Mellitus Treatment: A Randomized Clinical Trial. JAMA Surg. 2015;150(10):931–40.

10. Cummings DE, Arterburn DE, Westbrook EO, Kuzma JN, Stewart SD, Chan CP, et al. Gastric bypass surgery vs intensive lifestyle and medical intervention for type 2 diabetes: the CROSSROADS randomised controlled trial. Diabetologia. 2016;59(5):945–53.

11. Douros JD, Tong J, D’Alessio DA. The Effects of Bariatric Surgery on Islet Function, Insulin Secretion, and Glucose Control. Endocr Rev. 2019;40(5):1394–423.

12. Salehi M, Prigeon RL, D’Alessio DA. Gastric bypass surgery enhances glucagon-like peptide 1-stimulated postprandial insulin secretion in humans. Diabetes. 2011;60(9):2308–14.

13. Tian J, Huang S, Sun S, Ding L, Zhang E, Huang W. Bile acid signaling and bariatric surgery. Liver Res. 2017;1(4):208–13.

14. Klebanoff MJ, Corey KE, Chhatwal J, Kaplan LM, Chung RT, Hur C. Bariatric surgery for nonalcoholic steatohepatitis: A clinical and cost-effectiveness analysis. Hepatology. 2017;65(4):1156–64.

15. Ashrafian H, le Roux CW, Darzi A, Athanasiou T. Effects of bariatric surgery on cardiovascular function. Circulation. 2008;118(20):2091–102.

16. Akalestou E, Suba K, Lopez-Noriega L, Georgiadou E, Chabosseau P, Leclerc I, et al. Intravital imaging of islet Ca2+ dynamics reveals enhanced β cell connectivity after bariatric surgery in mice. bioRxiV. 2020.

17. Akalestou E, Lopez-Noriega L, Leclerc I, Rutter GA. Vertical sleeve gastrectomy lowers kidney SGLT2 expression in the mouse. bioRxiv. 2019.

18. Perez-Pevida B, Escalada J, Miras AD, Fruhbeck G. Mechanisms Underlying Type 2 Diabetes Remission After Metabolic Surgery. Front Endocrinol (Lausanne). 2019;10:641.

19. Nannipieri M, Mari A, Anselmino M, Baldi S, Barsotti E, Guarino D, et al. The role of beta-cell function and insulin sensitivity in the remission of type 2 diabetes after gastric bypass surgery. J Clin Endocrinol Metab. 2011;96(9):E1372–9.

20. Bornstein SR, Engeland WC, Ehrhart-Bornstein M, Herman JP. Dissociation of ACTH and glucocorticoids. Trends Endocrinol Metab. 2008;19(5):175–80.

21. Lamounier-Zepter V, Ehrhart-Bornstein M, Bornstein SR. Metabolic syndrome and the endocrine stress system. Horm Metab Res. 2006;38(7):437–41.

22. Klein NA, Andersen RN, Casson PR, Buster JE, Kramer RE. Mechanisms of insulin inhibition of ACTH-stimulated steroid secretion by cultured bovine adrenocortical cells. J Steroid Biochem Mol Biol. 1992;41(1):11–20.

23. Andreis PG, Malendowicz LK, Neri G, Tortorella C, Nussdorfer GG. Effects of glucagon and glucagon-like peptide-1 on glucocorticoid secretion of dispersed rat adrenocortical cells. Life Sci. 1999;64(24):2187–97.

24. Mazzocchi G, Gottardo L, Aragona F, Albertin G, Nussdorfer GG. Glucagon inhibits ACTH-stimulated cortisol secretion from dispersed human adrenocortical cells by activating unidentified receptors negatively coupled with the adenylate cyclase cascade. Horm Metab Res. 2000;32(7):265–8.

25. Kinzig KP, D’Alessio DA, Herman JP, Sakai RR, Vahl TP, Figueiredo HF, et al. CNS glucagon-like peptide-1 receptors mediate endocrine and anxiety responses to interoceptive and psychogenic stressors. J Neurosci. 2003;23(15):6163–70.

26. Gil-Lozano M, Perez-Tilve D, Alvarez-Crespo M, Martis A, Fernandez AM, Catalina PA, et al. GLP-1(7-36)-amide and Exendin-4 stimulate the HPA axis in rodents and humans. Endocrinology. 2010;151(6):2629–40.

27. Pullen TJ, Huising MO, Rutter GA. Analysis of Purified Pancreatic Islet Beta and Alpha Cell Transcriptomes Reveals 11beta-Hydroxysteroid Dehydrogenase (Hsd11b1) as a Novel Disallowed Gene. Front Genet. 2017;8:41.

28. Rebuffe-Scrive M, Bronnegard M, Nilsson A, Eldh J, Gustafsson JA, Bjorntorp P. Steroid hormone receptors in human adipose tissues. J Clin Endocrinol Metab. 1990;71(5):1215–9.

29. Dourakis SP, Sevastianos VA, Kaliopi P. Acute severe steatohepatitis related to prednisolone therapy. Am J Gastroenterol. 2002;97(4):1074–5.

30. Samuel VT, Petersen KF, Shulman GI. Lipid-induced insulin resistance: unravelling the mechanism. Lancet. 2010;375(9733):2267–77.

31. Dimitriadis G, Leighton B, Parry-Billings M, Sasson S, Young M, Krause U, et al. Effects of glucocorticoid excess on the sensitivity of glucose transport and metabolism to insulin in rat skeletal muscle. Biochem J. 1997;321 (Pt 3):707–12.

32. Livingstone DE, Jones GC, Smith K, Jamieson PM, Andrew R, Kenyon CJ, et al. Understanding the role of glucocorticoids in obesity: tissue-specific alterations of corticosterone metabolism in obese Zucker rats. Endocrinology. 2000;141(2):560–3.

33. Rask E, Olsson T, Soderberg S, Andrew R, Livingstone DE, Johnson O, et al. Tissue-specific dysregulation of cortisol metabolism in human obesity. J Clin Endocrinol Metab. 2001;86(3):1418–21.

34. Stomby A, Andrew R, Walker BR, Olsson T. Tissue-specific dysregulation of cortisol regeneration by 11betaHSD1 in obesity: has it promised too much? Diabetologia. 2014;57(6):1100–10.

35. Masuzaki H, Paterson J, Shinyama H, Morton NM, Mullins JJ, Seckl JR, et al. A transgenic model of visceral obesity and the metabolic syndrome. Science. 2001;294(5549):2166–70.

36. Rosenstock J, Banarer S, Fonseca VA, Inzucchi SE, Sun W, Yao W, et al. The 11-beta-hydroxysteroid dehydrogenase type 1 inhibitor INCB13739 improves hyperglycemia in patients with type 2 diabetes inadequately controlled by metformin monotherapy. Diabetes Care. 2010;33(7):1516–22.

37. Feig PU, Shah S, Hermanowski-Vosatka A, Plotkin D, Springer MS, Donahue S, et al. Effects of an 11beta-hydroxysteroid dehydrogenase type 1 inhibitor, MK-0916, in patients with type 2 diabetes mellitus and metabolic syndrome. Diabetes Obes Metab. 2011;13(6):498–504.

38. Woods CP, Corrigan M, Gathercole L, Taylor A, Hughes B, Gaoatswe G, et al. Tissue specific regulation of glucocorticoids in severe obesity and the response to significant weight loss following bariatric surgery (BARICORT). J Clin Endocrinol Metab. 2015;100(4):1434–44.

39. Simonyte K, Olsson T, Naslund I, Angelhed JE, Lonn L, Mattsson C, et al. Weight Loss after Gastric Bypass Surgery in Women Is Followed by a Metabolically Favorable Decrease in 11 beta-Hydroxysteroid Dehydrogenase 1 Expression in Subcutaneous Adipose Tissue. J Clin Endocr Metab. 2010;95(7):3527–31.

40. Methlie P, Dankel S, Myhra T, Christensen B, Gjerde J, Fadnes D, et al. Changes in adipose glucocorticoid metabolism before and after bariatric surgery assessed by direct hormone measurements. Obesity. 2013;21(12):2495–503.

41. Jorgensen NB, Dirksen C, Bojsen-Moller KN, Jacobsen SH, Worm D, Hansen DL, et al. Exaggerated glucagon-like peptide 1 response is important for improved beta-cell function and glucose tolerance after Roux-en-Y gastric bypass in patients with type 2 diabetes. Diabetes. 2013;62(9):3044–52.

42. Chambers AP, Smith EP, Begg DP, Grayson BE, Sisley S, Greer T, et al. Regulation of gastric emptying rate and its role in nutrient-induced GLP-1 secretion in rats after vertical sleeve gastrectomy. Am J Physiol Endocrinol Metab. 2014;306(4):E424–32.

43. Al-Sabah S, Alasfar F, Al-Khaledi G, Dinesh R, Al-Saleh M, Abul H. Incretin response to a standard test meal in a rat model of sleeve gastrectomy with diet-induced obesity. Obes Surg. 2014;24(1):95–101.

44. Ettleson MD, Lager CJ, Kraftson AT, Esfandiari NH, Oral EA. Roux-en-Y gastric bypass versus sleeve gastrectomy: risks and benefits. Minerva Chir. 2017;72(6):505–19.

45. Hutch CR, Sandoval D. The Role of GLP-1 in the Metabolic Success of Bariatric Surgery. Endocrinology. 2017;158(12):4139–51.

46. Murphy R, Clarke MG, Evennett NJ, John Robinson S, Lee Humphreys M, Hammodat H, et al. Laparoscopic Sleeve Gastrectomy Versus Banded Roux-en-Y Gastric Bypass for Diabetes and Obesity: a Prospective Randomised Double-Blind Trial. Obes Surg. 2018;28(2):293–302.

47. Garibay D, Lou J, Lee SA, Zaborska KE, Weissman MH, Sloma E, et al. beta Cell GLP-1R Signaling Alters alpha Cell Proglucagon Processing after Vertical Sleeve Gastrectomy in Mice. Cell Rep. 2018;23(4):967–73.

48. Douros JD, Niu J, Sdao S, Gregg T, Fisher-Wellman K, Bharadwaj M, et al. Sleeve gastrectomy rapidly enhances islet function independently of body weight. JCI Insight. 2019;4(6).

49. Larraufie P, Roberts GP, McGavigan AK, Kay RG, Li J, Leiter A, et al. Important Role of the GLP-1 Axis for Glucose Homeostasis after Bariatric Surgery. Cell Rep. 2019;26(6):1399–408 e6.

50. McGavigan AK, Garibay D, Henseler ZM, Chen J, Bettaieb A, Haj FG, et al. TGR5 contributes to glucoregulatory improvements after vertical sleeve gastrectomy in mice. Gut. 2017;66(2):226–34.

51. Gong S, Miao YL, Jiao GZ, Sun MJ, Li H, Lin J, et al. Dynamics and correlation of serum cortisol and corticosterone under different physiological or stressful conditions in mice. PLoS One. 2015;10(2):e0117503.

52. van der Velden VH. Glucocorticoids: mechanisms of action and anti-inflammatory potential in asthma. Mediators Inflamm. 1998;7(4):229–37.

53. Quaroni A, Tian JQ, Goke M, Podolsky DK. Glucocorticoids have pleiotropic effects on small intestinal crypt cells. Am J Physiol. 1999;277(5):G1027–40.

54. Akalestou E, Genser L, Rutter GA. Glucocorticoid Metabolism in Obesity and Following Weight Loss. Front Endocrinol (Lausanne). 2020;11:59.

55. Tomlinson JW, Finney J, Hughes BA, Hughes SV, Stewart PM. Reduced glucocorticoid production rate, decreased 5alpha-reductase activity, and adipose tissue insulin sensitization after weight loss. Diabetes. 2008;57(6):1536–43.

56. Purnell JQ, Kahn SE, Samuels MH, Brandon D, Loriaux DL, Brunzell JD. Enhanced cortisol production rates, free cortisol, and 11beta-HSD-1 expression correlate with visceral fat and insulin resistance in men: effect of weight loss. Am J Physiol Endocrinol Metab. 2009;296(2):E351–7.

57. Johnstone AM, Faber P, Andrew R, Gibney ER, Elia M, Lobley G, et al. Influence of short-term dietary weight loss on cortisol secretion and metabolism in obese men. Eur J Endocrinol. 2004;150(2):185–94.

58. Morrow J, Gluck M, Lorence M, Flancbaum L, Geliebter A. Night eating status and influence on body weight, body image, hunger, and cortisol pre- and post-Roux-en-Y Gastric Bypass (RYGB) surgery. Eat Weight Disord. 2008;13(4):e96–9.

59. Larsen JK, van Ramshorst B, van Doornen LJ, Geenen R. Salivary cortisol and binge eating disorder in obese women after surgery for morbid obesity. Int J Behav Med. 2009;16(4):311–5.

60. Ruiz-Tovar J, Oller I, Galindo I, Llavero C, Arroyo A, Calero A, et al. Change in levels of C-reactive protein (CRP) and serum cortisol in morbidly obese patients after laparoscopic sleeve gastrectomy. Obes Surg. 2013;23(6):764–9.

61. Valentine AR, Raff H, Liu H, Ballesteros M, Rose JM, Jossart GH, et al. Salivary cortisol increases after bariatric surgery in women. Horm Metab Res. 2011;43(8):587–90.

62. Argaud D, Zhang Q, Pan W, Maitra S, Pilkis SJ, Lange AJ. Regulation of rat liver glucose-6-phosphatase gene expression in different nutritional and hormonal states: gene structure and 5’-flanking sequence. Diabetes. 1996;45(11):1563–71.

63. Xu C, He J, Jiang H, Zu L, Zhai W, Pu S, et al. Direct effect of glucocorticoids on lipolysis in adipocytes. Mol Endocrinol. 2009;23(8):1161–70.

64. Paterson JM, Morton NM, Fievet C, Kenyon CJ, Holmes MC, Staels B, et al. Metabolic syndrome without obesity: Hepatic overexpression of 11beta-hydroxysteroid dehydrogenase type 1 in transgenic mice. Proc Natl Acad Sci U S A. 2004;101(18):7088–93.

65. Holst JJ. The physiology of glucagon-like peptide 1. Physiol Rev. 2007;87(4):1409–39.

66. Beato M. Gene regulation by steroid hormones. Cell. 1989;56(3):335–44.

67. Opherk C, Tronche F, Kellendonk C, Kohlmuller D, Schulze A, Schmid W, et al. Inactivation of the glucocorticoid receptor in hepatocytes leads to fasting hypoglycemia and ameliorates hyperglycemia in streptozotocin-induced diabetes mellitus. Mol Endocrinol. 2004;18(6):1346–53.

68. Shen Y, Roh HC, Kumari M, Rosen ED. Adipocyte glucocorticoid receptor is important in lipolysis and insulin resistance due to exogenous steroids, but not insulin resistance caused by high fat feeding. Mol Metab. 2017;6(10):1150–60.

69. Grayson BE, Hakala-Finch AP, Kekulawala M, Laub H, Egan AE, Ressler IB, et al. Weight loss by calorie restriction versus bariatric surgery differentially regulates the hypothalamo-pituitary-adrenocortical axis in male rats. Stress. 2014;17(6):484–93.

70. Rask E, Simonyte K, Lonn L, Axelson M. Cortisol metabolism after weight loss: associations with 11 beta-HSD type 1 and markers of obesity in women. Clin Endocrinol (Oxf). 2013;78(5):700–5.

71. Sumner SM, Regier PJ, Case JB, Ellison GW. Evaluation of suture reinforcement for stapled intestinal anastomoses: 77 dogs (2008-2018). Vet Surg. 2019;48(7):1188–93.

